# Does heat tolerance actually predict animals’ geographic thermal limits?

**DOI:** 10.1101/2021.11.30.468083

**Authors:** Agustín Camacho, Miguel Trefaut Rodrigues, Refat Jayyusi, Mohamed Harun, Marco Geraci, Catarina Vinagre, Miguel Carretero, Miguel Tejedo

## Abstract

To understand species′ climatic vulnerability, our measures of species’ thermal tolerance should predict their geographic thermal limits. Yet, this assumption is ungranted. We tested if animals′ heat tolerance restrict the warmest temperatures they can live at (Tmax), distinguishing among species differently challenged by their thermal environment. For that, we compiled 2350 measurements of species’ heat tolerance indexes and corresponding Tmax, measured at different microhabitats. We show that reptiles, a flagship for climatic vulnerability studies, are particularly unbounded by their heat tolerance. Contrarily, tolerance restricted marine fish’ geography in a non-linear fashion which contrasts with terrestrial taxa. Behavioral tolerance indexes, widely used to predict vulnerability, predicted Tmax inconsistently across Tmax indexes, or were inversely related to it. Animals’ heat tolerance restricts geographic limits more strongly for more thermally challenged species. In turn, factors uncoupling heat tolerance and Tmax (plasticity, thermoregulation, adaptation) should be more important for less thermally challenged species.

**Significance Statement:** To identify climatic vulnerability, heat tolerance indices need to predict species′ thermal limits to geographic distribution. Yet, we show that heat tolerance predicts geographic limits quite heterogeneously, depending on taxa, the type of measure of heat tolerance and how challenged are species at their hottest known location. Particularly, reptiles, a flagship of vulnerability studies, were less bounded by tolerance than taxa regarded as more capable to evade high temperatures, due to their capacity to evaporate water, find refuge, or migrate (Ex. amphibians, arthropods, birds and mammals). Measures of species’ behavioral heat tolerance may still need to improve. Factors thought to uncouple thermal tolerance and geographic limits should be stronger for less thermally challenged species.

## Main

A reliable identification of the climatic vulnerability of populations is crucial to correctly allocate the huge investments necessary to preserve biodiversity from climate warming^1^. Previous evaluations have used different thermal tolerance indexes, searching for global patterns in species’ vulnerability^2,3,4,5,6,7^, that rely on a key assumption.

This is that heat tolerance restricts the sites supporting individual’s survival, dispersal, or population’s growth, to the level of stablishing the warm edges of species’ geographic distribution^8^. This assumption, is however ungranted for most animal groups, since they readily decouple their body temperatures from exceedingly hot temperatures across their ranges^6,9,10^, potentially buffering the evolution of heat tolerance itself^11,12^. Several other factors may also unlink species’ geographic limits from their known measures of heat tolerance. For example, local adaptation at species’ warm edges, due to exposure to relatively extreme temperatures^13^, plasticity and adaptation in heat tolerance traits^14,15^, species interactions blocking dispersal^16,17^, incorrect estimation of heat tolerance^18,19^ and environmental temperatures^20^, or the wallacean shortfall.

Correctly predicting temperatures that limit species’ distribution is imperative to identify the conditions that set species’ geographic warm edges, where further climatic warming should make species’ populations’ dwindle^21,22^. However, previous global patterns of geographic variation in thermal tolerance and climatic risk have compared species’ heat tolerance with the thermal environment of the sampling sites of the measured individuals^6,21,23^. This conservative approach helps avoiding the outlined decoupling factors. However, it means we still do not know whether heat tolerance, insofar measured in animals, widely predicts the environmental temperatures that restrict species’ geographic distribution, or how it does so for most groups.

We tested whether species’ thermal tolerance predicts maximum environmental temperatures at each species’ known warm edges during the hottest time of the year (hereafter Tmax, see methods). Both, tolerance and Tmax have been estimated using many indexes, often associated to specific taxa (see methods). Different tolerance indexes describe different levels of thermal stress for organisms (ex. acute to chronic^24^). Thus, we assorted these indexes within three main groups that induce declining levels of thermal stress (Thermal limits, upper limits to optimal physiological temperatures, and indexes of behavioral tolerance, see methods), and taxa. We used quantile mixed models to separate species according to how much Tmax would challenge their heat tolerance. Thus, higher quantiles represent species that reach higher Tmax at their geographic warm edges, given their heat tolerance levels.

Our first group of heat tolerance indexes, the physiological thermal limits (hereafter CTmax, from Critical Thermal Maximum), represent temperatures that can kill during acute exposures (i.e. minutes, See methods). The CTmax does restrict the highest Tmax values (i.e., significantly predicts its 90^th^ percentile) across marine fish, and terrestrial ectotherms, with the exception of reptiles (Fig.1, Figs. S2: A B C, Table 1). While results for Marine fish are robust across Tmax indexes (Ocean’s surface and mid depth temperatures), the relationships of terrestrial ectotherms varied more across Tmaxexp, Tmaxprot and Tmaxair, estimated at exposed and protected microhabitats, and in the air, respectively (Tables S1: A, B, C; Figs S2: A, B, C, See methods). Thus, CTmax can be used to spot species’ climatic limits for the distribution of ectotherms from different realms, but with special caution for terrestrial species as Tmax may be unrelated to CTmax. This is particularly true for reptiles, a flagship group for the climatic vulnerability problem, whose CTmax was less strongly or not related to Tmax (Table 1).

**Table 1.**
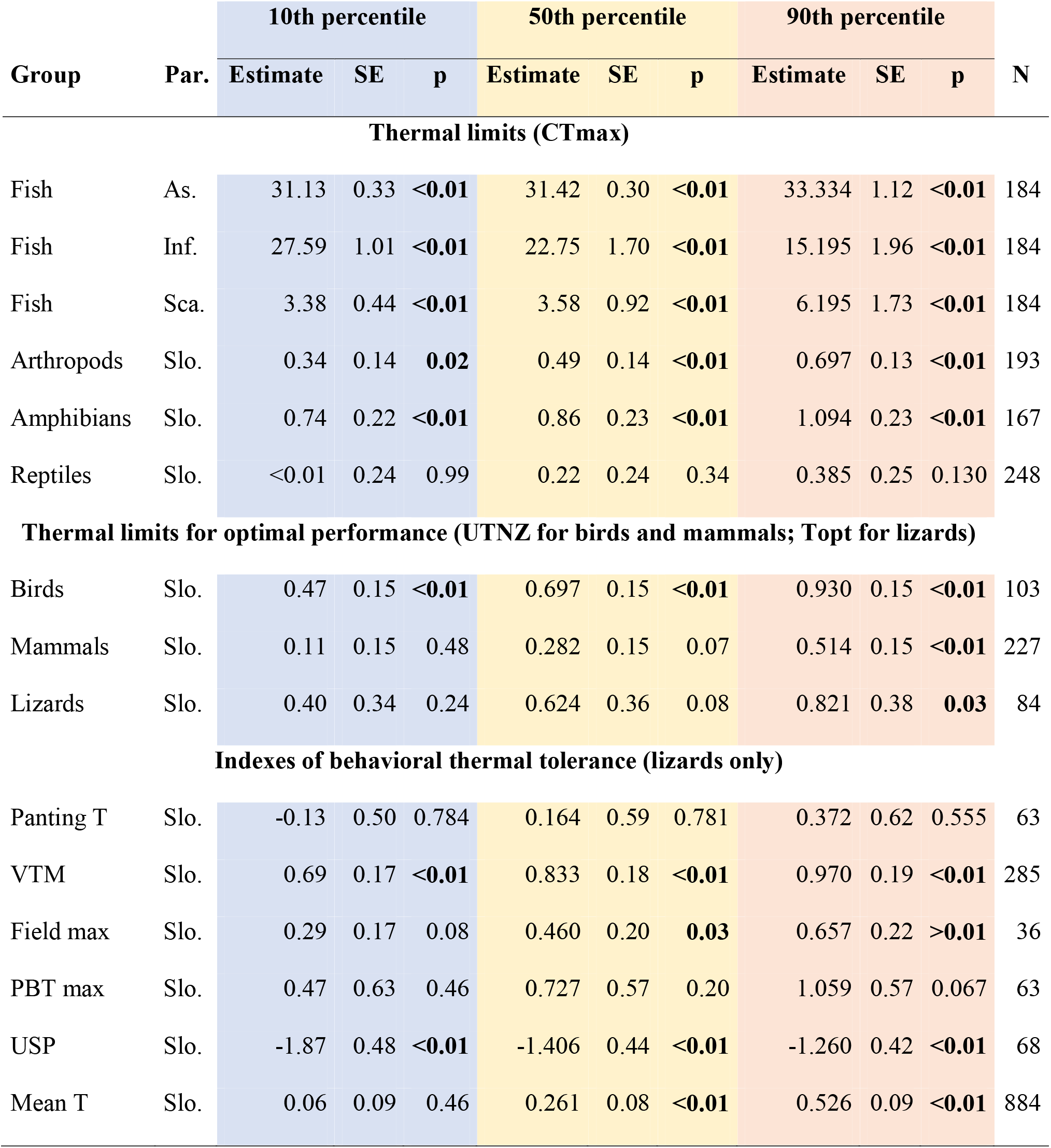
Relationships between Tmax and Heat tolerance across animals. Describes parameters, standard errors (SE) and p-values from nonlinear (marine fish) and linear (other animal groups) quantile mixed models for the 10^th^, 50^th^, and 90th percentiles of Tmax conditional on índices of thermal tolerance. CTmax, critical thermal maximum; UTNZ: upper thermoneutral zone limit; Topt: optimal temperature for sprint speed; Panting T: body temperature that induces panting; VTM: voluntary thermal maximum, body temperature that induces retreat; Field max: maximum temperature observed in the field; PBT max: maximum body temperature measured in a laboratory thermal gradient; USP: Upper Set Point 75^th^ percentile of preferred temperatures, ; Mean T: mean body temperature of active individuals; Par.=Parameter, As.=Asymptote, Inf.=Inflection point, Sca.=Scale, Slo.=Slope, <N=species number per analysis. See methods for definitions of tolerance indexes. Correlations’ intercepts can be observed in Table S1. Colors relate to quantiles in Fig. 1. Tmaxexp source: Microclim for terrestrial spp./Bio-oracle for marine ones.

| Group | Par. | 10th percentile |  |  | 50th percentile |  |  | 90th percentile |  |  | N |
| --- | --- | --- | --- | --- | --- | --- | --- | --- | --- | --- | --- |
|  |  | Estimate | SE | p | Estimate | SE | p | Estimate | SE | p |  |
| Thermal limits (CTmax) |  |  |  |  |  |  |  |  |  |  |  |
| Fish | As. | 31.13 | 0.33 | <0.01 | 31.42 | 0.30 | <0.01 | 33.334 | 1.12 | <0.01 | 184 |
| Fish | Inf. | 27.59 | 1.01 | <0.01 | 22.75 | 1.70 | <0.01 | 15.195 | 1.96 | <0.01 | 184 |
| Fish | Sca. | 3.38 | 0.44 | <0.01 | 3.58 | 0.92 | <0.01 | 6.195 | 1.73 | <0.01 | 184 |
| Arthropods | Slo. | 0.34 | 0.14 | 0.02 | 0.49 | 0.14 | <0.01 | 0.697 | 0.13 | <0.01 | 193 |
| Amphibians | Slo. | 0.74 | 0.22 | <0.01 | 0.86 | 0.23 | <0.01 | 1.094 | 0.23 | <0.01 | 167 |
| Reptiles | Slo. | <0.01 | 0.24 | 0.99 | 0.22 | 0.24 | 0.34 | 0.385 | 0.25 | 0.130 | 248 |
| Thermal limits for optimal performance (UTNZ for birds and mammals; Topt for lizards) |  |  |  |  |  |  |  |  |  |  |  |
| Birds | Slo. | 0.47 | 0.15 | <0.01 | 0.697 | 0.15 | <0.01 | 0.930 | 0.15 | <0.01 | 103 |
| Mammals | Slo. | 0.11 | 0.15 | 0.48 | 0.282 | 0.15 | 0.07 | 0.514 | 0.15 | <0.01 | 227 |
| Lizards | Slo. | 0.40 | 0.34 | 0.24 | 0.624 | 0.36 | 0.08 | 0.821 | 0.38 | 0.03 | 84 |
| Indexes of behavioral thermal tolerance (lizards only) |  |  |  |  |  |  |  |  |  |  |  |
| Panting T | Slo. | -0.13 | 0.50 | 0.784 | 0.164 | 0.59 | 0.781 | 0.372 | 0.62 | 0.555 | 63 |
| VTM | Slo. | 0.69 | 0.17 | <0.01 | 0.833 | 0.18 | <0.01 | 0.970 | 0.19 | <0.01 | 285 |
| Field max | Slo. | 0.29 | 0.17 | 0.08 | 0.460 | 0.20 | 0.03 | 0.657 | 0.22 | >0.01 | 36 |
| PBT max | Slo. | 0.47 | 0.63 | 0.46 | 0.727 | 0.57 | 0.20 | 1.059 | 0.57 | 0.067 | 63 |
| USP | Slo. | -1.87 | 0.48 | <0.01 | -1.406 | 0.44 | <0.01 | -1.260 | 0.42 | <0.01 | 68 |
| Mean T | Slo. | 0.06 | 0.09 | 0.46 | 0.261 | 0.08 | <0.01 | 0.526 | 0.09 | <0.01 | 884 |

Studies comparing species’ heat tolerance and the climate of collection sites have portraited arthropods and amphibians as especially able to decouple their body temperatures from high environmental temperatures (ex. hiding in small cracks or evaporating body water^6,21^). Yet, their heat tolerance predicts their geographic thermal limits more strongly than reptiles’ heat tolerance. Compared to the former, reptiles exhibit similar CTmax acclimation capacities^25^, and their geographic distribution seems not contrastingly solved. Also, studied arthropods show less constrained CTmax evolution than Squamates^26^, being able to change their CTmax more than two degrees in three generations. In turn, reptiles do have lower body water loss rates^27^ and often bigger size, compared to amphibians and arthropods, respectively. The large variation shown in water loss by this group^28^ should help identifying the importance of this trait for CTmax-distribution relationships.

Biotic interactions may limit range more strongly at warmer edges, compared to cold edges^44^. We did not account for biotic interactions in our analysis and thus it is either possible that 1) biotic interactions contribute to blur tolerance-Tmax relationships at less challenging warmer edges, 2) intensify the importance of a constitutively high thermal tolerance at more challenging warmer edges, or 3) there is no interaction between these factors^17^.

Accordingly, there should be a feedback between Tmax at warm edges and thermal tolerance, where encountering more challenging Tmax may select for higher heat tolerance, and it in turn may help the expansion of warm distribution edges. In a context of climatic warming, warm edges can be either the trailing fronts of thermally challenged species or the advance fronts for not so challenged ones. Species more challenged by the new Tmax conditions may need to evolve higher heat tolerance. In turn, at less challenging ones, a plastic heat tolerance might be more useful.

The strong restriction of CTmax on geographic thermal limits of marine fish (Fig.1, Table 1) agrees with their capability to full fill their thermal niches across latitudinal ranges. The CTmax of warm-adapted fish (CTmax > 35 °C) surpasses sea surface’s thermal maxima registered in geographic datasets (~35 °C, Sbrocco & Barber 2013 29). This and the very low CTmax of polar fish^30^, strongly restricting Tmax, leads to strong and non-linear relationships between CTmax and Tmax for this group (Fig. 1, Supporting figures, Supporting tables set 1). CTmax levels over maximum sea Tmax should facilitate fish dispersal across equatorial coasts, where the sea is warmest^21^. This physiological liberation helps explaining why fish species distributed through the equator often have larger geographic ranges^31^. The highest CTmax among marine fish likely have evolved in response to selective pressures existing at globally hyperthermic microhabitats (ex. coastal rock pools exceeding 41 °C^32^).

**Figure 1.**
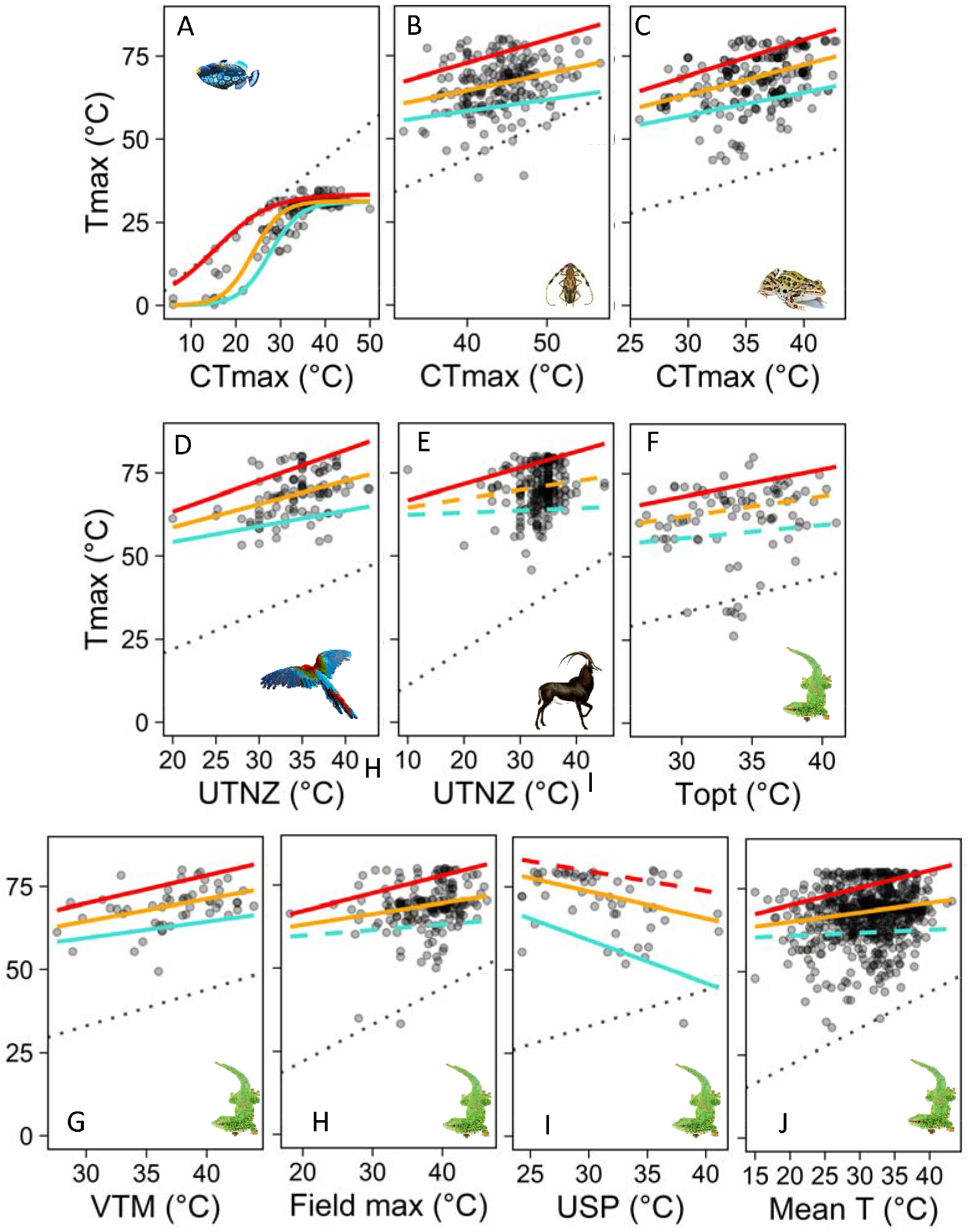
Upper thermal tolerance indexes capable of predicting Geographic thermal limits (Tmax: Maximum temperature available) among different animal groups. A-C) Thermal limits as measured in marine fish, terrestrial arthropods, amphibians, and terrestrial reptiles. E-G) Upper limits of physiologically optimal temperatures measured in birds, mammals, and lizards. H-K) Indexes of behavioral thermal tolerance measured in reptiles (see definitions in Table 1 and methods). Dotted line represents a 1:1 relationship between thermal tolerance and the geographic thermal limit. Solid lines represent the slopes of phylogenetically detrended predictions of Tmax based on thermal physiology. Red, orange and blue lines represent predicted 90th, 50th, and 10th conditional percentiles of Tmax on thermal tolerance, respectively. Slashed lines represent non-significant associations. See graphs for all heat tolerance and Tmax indexes in Fig. S2.

Our second group of tolerance indexes identify thermal levels over which physiological maintenance costs become especially high. Long exposures to higher temperatures undermine sustained performance, energy assimilation and hydration and can be ultimately lethal (see methods). Among endotherms, these levels have been more often measured as the upper limit of the thermoneutral zone (UTNZ, see methods), while in lizards, they have been often represented by the body temperatures for fastest sprint speed (Topt, See methods). In agreement with Khaliq et al^4^ endotherms’ warm edges are generally predicted by their UTNZ. Yet, again, squamates appear again as less, or no geographically restricted by their optimal temperatures (see Figs. 1, and S2 A, B, C, D, and Tables S1). Apart from the factors mentioned in the introduction, endotherms might be more powerful dispersers than lizards, but still blocked by too hot temperatures. Also, hiding from exceedingly high temperatures (ex. aestivating^33^) is less costly for lizards than for endotherms, suggesting they rely more on heat tolerance for extending their ranges than lizards, this seems to be particularly true for birds, less able to use underground refuges.

Behavioral indexes, estimated for different ranges of temperatures voluntarily tolerated by animals, could provide less stressful, more comparable, and readily measurable indexes of heat tolerance for endotherms and ectotherms (ex. Panting^34^). We made a compilation of these indexes for reptile species (mostly lizards), where they have been most extensively used to assess climatic vulnerability^3,9^. Specifically, we obtained data on species’ panting temperatures, voluntary thermal maxima, maximum preferred temperatures, maximum field temperatures, and both, the upper set point and the mean of body temperatures recorded for active individuals (see methods). However, behavioral traits predicted Tmax quite heterogeneously among tolerance and Tmax indexes (See below, and Tables S1 A, B, C, D). The weakest predictors were panting and the maximum of preferred temperatures. Panting costs much body water, thus chronic use of this behavior can lead to progressive decrement in body condition^35^. Even drought-adapted species are reluctant to pant when dehydrated^34^. Thus, this behavior might only help to avoid instant heat shock, rather than to survive stressfully hot, and often dry, periods at warm edges. Instead, measures of maximal preferred temperatures traditionally carry over many methodological pitfalls^18^, affecting their use for this end.

Instead, the most robust predictors of Tmax where the maximum field body temperatures and the mean preferred body temperatures. The latter shows a blurry relationship with Tmax (Fig. 1J) and small slopes compared to other behavioral indexes (Fig. 3). Thus, the significant correlations are likely favored by the much larger data availability. Still, the former might provide particularly useful data. Lizards adjust their activity levels and active body temperatures with respect to limiting factors, such like hydration state^36^. Thus, if collected in a controlled fashion (for animals under environmental distress), maximum field body temperatures might integrate these factors and show which temperatures actually limit species’ activity and geographic warm edges. The case of the upper set point is also interesting. It negatively correlates with Tmax (Fig. 1, Table 1, and Tables S1, A, B, C, D) which makes it less useful to predict climatic vulnerability. Since mean body temperatures showed blurry but always positive relations with Tmax, lizards living at particularly cold distribution edges might raise their upper set point. Doing this should boost their growth rates^37^ and locomotor function^38^ within still tolerable thermal levels. Our results highlight the need of field evaluations of restrictions predicted by behavioral indexes, which are so far rare^3,39^.

**Figure 2.**
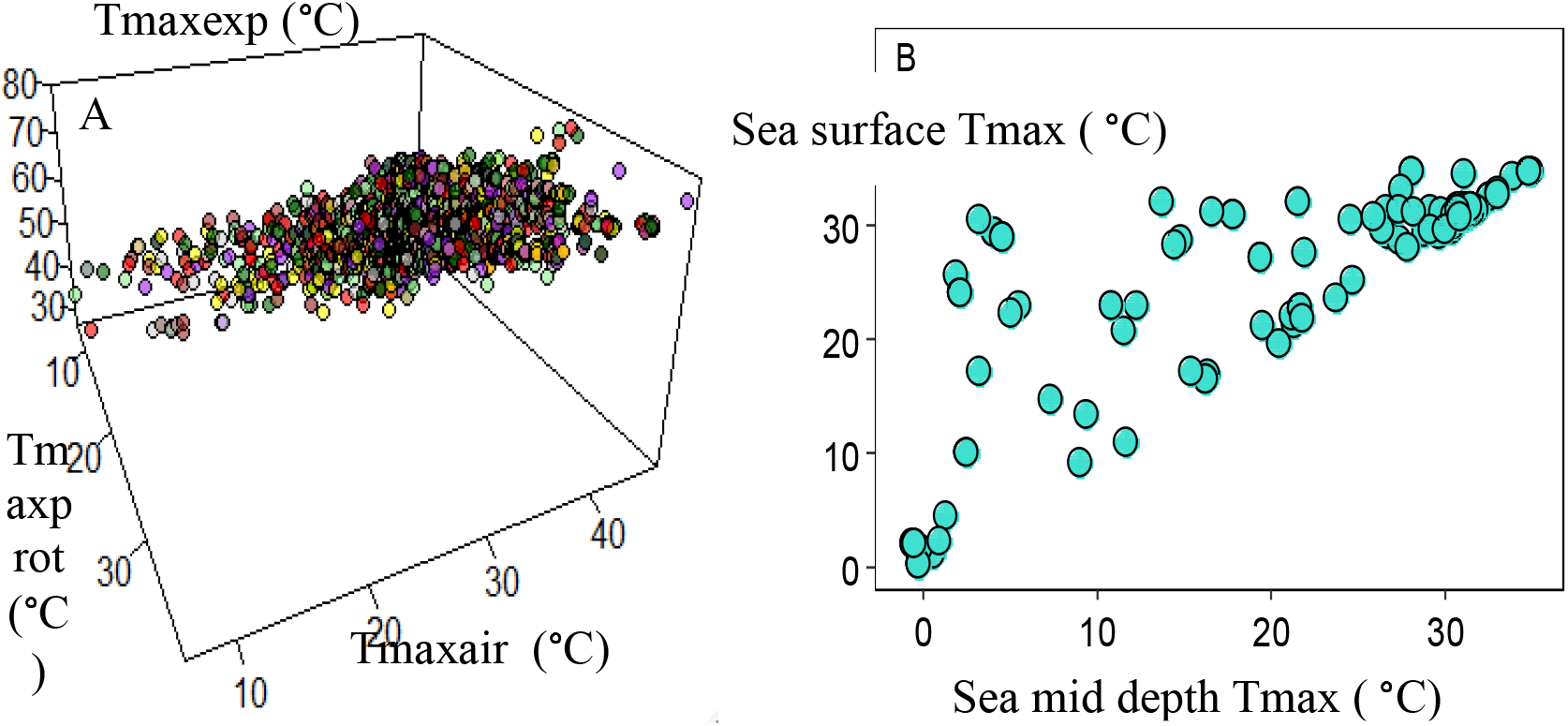
Relationships among Tmax measured at different microhabitats of species’ geographic warm edges. A panel shows relationships for terrestrial animals. B panel shows relationships for Marine fish. Whereas relationships for terrestrial species are strongly linear, in the sea, differences between mid and surface Tmax change across increasingly hot geographic thermal limits.

**Figure 3.**
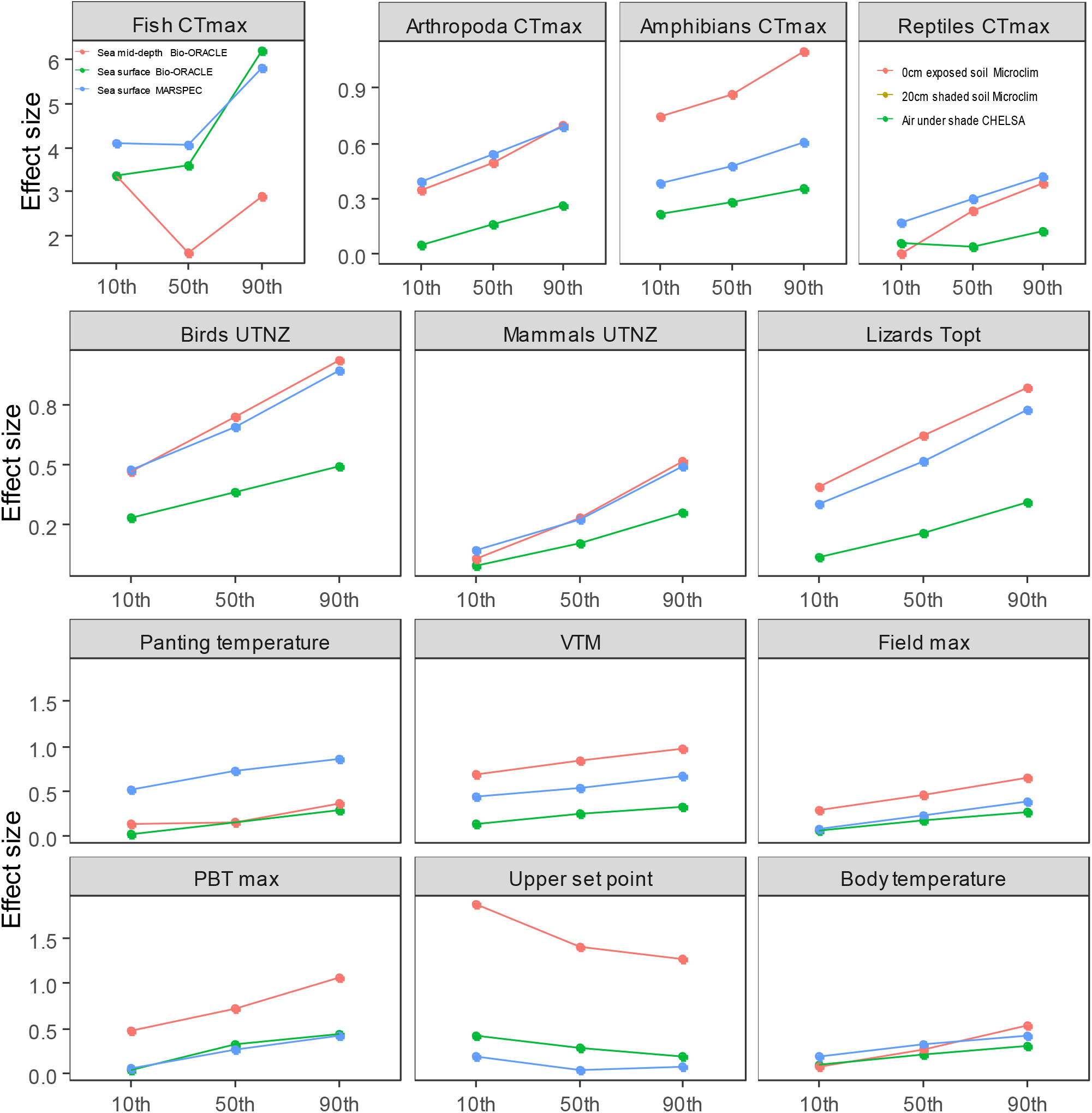
Effect sizes of relationships between thermal tolerance on the geographic thermal limit for species differently challenged by temperatures at their geographic thermal limit. Except for marine fish and the upper set point of lizards, thermal tolerance is more strongly linked to the geographic thermal limit of species reaching places with higher environmental temperatures with respect to their thermal tolerance (species at the 90^th^ percentile, see methods). See Table 1 legend for definition of indexes, with three different measures of maximal temperatures (Tmax protected, Tmax exposed and Tmax air). In the case of marine fish, instead of Tmaxair, we included a different estimate of Tmax exposed, see methods). See supporting tables set 2 for tests’ results.

Among thermal tolerance indexes, Tmaxexp exhibited stronger correlations with thermal tolerance, than Tmaxprot and Tmaxair. Steeper slopes are expected for Tmaxexp, given that its range of variation is much wider than Tmaxprot and Tmaxair (Ranges: Texp: 53.8, 26.2-80; Tprot: 31.8, 6.4-38.2 Tair: 40.8, 6.4-47.2). These temperatures may vary differently with latitude, with Tmax prot being usually higher at tropical regions^6^. However, when sampled across species’ warm edges, they are linearly and strongly correlated with each other across the warm edges of terrestrial species (Fig. 3 A, Tables S2 A, B, C). Such covariation translates into varied sources of thermal stress for terrestrial species at their geographic limits. For example, the more environmental temperatures at exposed sites depart from thermal tolerance, the more body heating rates should speed up^40^. This should increase overheating risks for small ectotherms when crossing exposed microhabitats during their daily activities^41^. Likewise, at geographic warm edges of terrestrial species, Tmaxair, measured in the shade, often reach levels close to CTmax (particularly for species of the 90^th^ percentile). This means that individuals will likely experience sublethal temperatures for some periods even in the shade. In such situations, longer resistance to sublethal thermal levels should be particularly adaptive^19^. Finally, since Tmaxprot, measured at underground shelters lay well below the thermal tolerance of organisms (Fig. S2C; Table S2C), we propose that these shelters are often unavailable (Ex. For birds) or ineffective to extend species’ ranges. Thus, shelters might be less useful as previously thought to protect them from warming-induced range contractions. When hidden for long periods, some species might decrease energy intake and mating rates^3^, limiting the utility of shelters for population persistence at warm edges. Differently from land, at the sea, Tmaxsurf and Tmaxmid correlate with great heteroscedasticity (Fig. 2B). Vertical thermoclines are smallest at the hottest and at the coldest sites (tropical and polar coasts), and largest at deeper seas. This situation hampers thermoregulation in these regions, making it more necessary to evolve a suitable CTmax to cross-by. In turn, at regions with steeper thermoclines in maximum temperatures, warm-adapted fish could extend their geographic warm edges via superficial and warmer waters, while cold adapted ones should use deeper layers to disperse.

Our global evidence (Fig.3, Table S2C) supports simulations predicting that CTmax might evolve more readily when species’ physiology are more strongly hit by thermal extremes^13^. When the necessary data becomes available to account for plasticity, our discovered relationships might somewhat change, as the relationships between thermal tolerance and latitudinal ranges^23^. However, such changes should be minor for the dispersal or persistence of species at their geographic warm edges, or otherwise it would have been unlikely to find so pervasive effects for the more challenged species. These results are further supported by the fact that warm-tolerant animals living at high environmental temperatures show less plastic CTmax^42^. Despite heat tolerance is widely viewed as relatively insensitive to thermal gradients^43^, it can covary strongly with challenging thermal extremes (Table 1, supporting tables set 1). In these situations, the strong interspecific covariation likely means that local adaptation is less capable of inducing geographic variation in heat traits (ex. between measured populations and populations at warm edges). Likewise, other restrictors to the evolvability of heat tolerance^26^ should be less important for more heat challenged species, too. Particularly, behaviors that buffer exposures to thermal extremes (The Bogert Effect^12^).

Behavior and plasticity (i.e. acclimation) make use of spatiotemporal heterogeneity in temperatures^23^, that may be relatively small at the hottest sites (ex. Fig. 2B; Sunday et al^6^). Specially given that temperatures are strongly correlated across microhabitats (Fig. 2B) at these edges. Thus, these traits might be less able to affect Tmax-Heat tolerance relationships at challenging warm edges. In agreement, heat tolerance-Tmax correlations were almost always weaker for species less challenged at their warm edges. The exceptions were two: First, the lizards’ Upper set point showed stronger correlations at colder sites (Fig. 3), further supporting the idea of behavioral adjustments being more important at relatively cold sites (see above). The second exception was for Tmax at Ocean′s mid depths, which shows no clear pattern (Fig. 3). Tmax indexes coming from high resolution climatic datasets (CHELSA and MARSPEC) behaved in parallel with Tmax indexes coming from 5 min datasets (Microclim and Biooracle, Fig. 3). This indicates that our findings are robust to the scale of the climatic datasets.

## Methods

### Estimation of Tmax for each species

Our dependent variable, Tmax, represents the maximum environmental temperature that each species will face off across its geographic distribution. In nature, Tmax may depend on each species’ ecology and ability to thermoregulate^1^. Because it is unknown which temperature most affects each species, we repeated the analyses over different Tmax estimates. For land species, we used: 1) Tmaxair, the temperature of shaded air at around 2m off the ground, as widely used in previous studies^2,3^, 2) Tmax protected (Tmaxpro), estimated for 20 cm underground under fully shaded soil, and 3) Tmaxexp, describing the temperature of the surface of soil fully exposed to sun rays. For marine species, Tmaxsurf represent temperatures measured at the sea surface, and Tmax mid depth represents the temperature at the middle of the water column.

For each species, we estimated Tmax from global raster layers describing current climate. Tmax is the mean of maximum temperatures of the hottest month, measured across a period of 20 recently past years. This method for summarizing extreme temperatures is known as bioclim 5^4^. Estimates of climatic vulnerability may vary with the resolution of the climatic layer used^5^. Therefore, we used the highest resolution available for each type of Tmax estimation. Pursuing this aim, for on land Tmax, we extracted Tmaxair from the CHELSA dataset (30 sec resolution). In turn, Tmaxpro and Tmaxexp were extracted from the microclim dataset (soil substrate, 5 min resolution^6^). For marine Tmax, we extracted sea surface from the MARSPEC dataset (30 sec resolution, MARSPEC^29^, and also, Tmax from sea surface and at mid depth from the Bioracle dataset (5 min resolution^7^). Since we are using datasets with different scales, different trends would be expected for their associated correlations. However, as we discuss in the main text, we found quite the opposite: datasets with different scale behaved similarly.

To estimate the Tmax of each species’ hottest location, we extracted these data for each location of each species and kept the maximum value. This temperature is hereafter used as the geographic thermal limit, and the location defined as the warm edge (AKA: trailing edge^8,9,10^. We obtained all known locations for each species with thermal tolerance data from the Global Biodiversity Information Facility (GIBF). We only used locations associated to specimens deposited in scientific collections. These records were cleaned for likely captivity sites (ex. zoos), outliers, lon/lat zeroed records, mirrored records, records in the sea for terrestrial species, and vice versa. We then obtained the maximum Tmax across all records of each species and its corresponding Tmax value (see supporting material-DATA). In the main text, we show results for Tmaxexp and Tmaxsurf (temperatures for exposed soil and sea surface), following recommendations by Sunday et al^11^, p4, 2nd paragraph), but results for other indices can be found in the supporting tables set 1 and in the supporting figures file. Besides, we discuss, in the main text, the robustness of our findings to using different Tmax indexes.

### Indexes of thermal tolerance

Our compiled thermal tolerance indexes represent a continuous gradient of thermal stress intensity, namely, from an acute extreme to a chronic medium heat stress, (^12^, pp: 331). Thus, we grouped thermal tolerance indexes accordingly, in three groups. Our first group represents physiological thermal limits. These are temperatures that kill almost immediately. For terrestrial ectotherms, they have been measured as the temperature that blocks individuals’ locomotion (CTMAX, from Critical Thermal Maximum^13^). We obtained them for 200 terrestrial arthropods, 151 adult amphibians, and 300 reptiles (Squamates and tortoises). For marine fish, we obtained data on the loss of equilibrium (LOE, 121 species), and of the lethal limit (63 species), namely, the mean of the first temperature that causes death in a fish during experimental heating^14^.

Our second group of indexes coarsely represent the upper thermal limits for optimal physiological performance. In lizards, these limits have been often measured as the body temperature that maximizes sprint speed (Topt^3^, 85 species in our dataset). Since, many physiological processes exhibit lower thermal optima, lizards tend to select temperatures below Topt, and even their sprint speed sharply decreases above that temperature^15^. In turn, among endotherms, these limits correspond to the Upper Thermoneutral Zone limits (UTNZ), an environmental temperature over which metabolic and water loss costs sharply increase^16^. We used measures for 105 birds and 232 mammals^17^.

Our third group of indexes describes body temperatures voluntarily tolerated by lizards, a widely studied organism in thermal physiology. For lizards, upper voluntary thermal limits are often represented by either the display of panting behavior^18^ (herein called Panting temperatures, 63 species), or the maximum body temperature measured in active individuals of a species. This latter index has been either measured in the field (herein called Field max, 285 species^19,20^, in laboratory thermal gradients (PBT max, for maximum preferred body temperatures, 36 species), or in heating chambers (Voluntary Thermal Maximum (VTM), 63 species). Others have used either the 75^th^ percentile (68 species^21,22^), or the mean (884 species^23^) of body temperatures, measured on active lizards. Also, to avoid misrepresentation of voluntary limits in field maximum temperatures, we excerpted field maximum temperatures measured in less than 10 individuals per species.

Most of these datasets use previous compilations for adult animals by various studies^3,11,14,17,18,20,22,23,24,25^. Further, we added new data whenever possible^26,27,28^ (more on the supporting material-data).

We also added data from a field trip to Mozambique that includes new data for the VTmax of 28 lizard species (see supporting methods file for details). Creating a fully comprehensive and up-to-date dataset is not the aim of this study. That would be very difficult because new data appear frequently and each species follows a long process for gathering and checking the locality records, obtain the three different Tmax estimates, and then re-run the analyses. Raw data and sources can be accessed at the supporting Data-tables. All these indexes have been measured in sites that may strongly differ in climatic conditions from each species’ warm edges. Precisely, the purpose of this study is to evaluate whether these indexes, in the way they have been measured, can still predict the climatic conditions at species’ warm edges.

Previous studies have tried to correct tolerance levels for effects of acclimation temperature, ramping rates or exposure duration^29^. Geographical patterns in thermal limits appears to be not strongly affected by covariates of acclimation temperature and ramping rate or exposure duration^29^. Besides, estimates for these effects are very rare and species’ specific^30,31^, which limits the effectivity of corrections for whole groups from data on a few species. Since acclimation is one of the multiple confusion factors discussed in the main text, the cases in which heat tolerance predicted Tmax less successfully (ex. among reptiles, or for lower quantiles), might indicate where future corrections like the these might provide more advance in the future.

### Analyses

#### Quantile correlations between Tmax and heat tolerance indexes

The relationship between the 10th, 50th, and 90th percentiles of the geographic thermal limit distribution and thermal tolerance was modelled via linear quantile mixed models (LQMM) [1] separately for all animal groups, except for fishes. For fishes only, a logistic nonlinear quantile mixed model (NLQMM) [2] was used instead. (Results in Table 1 and Figure 1).

The random effects specifications for LQMM, NLQMM, and AQMM were at the genus level. Standard errors and 95% confidence intervals were obtained via bootstrap with 199 replications. The analysis was conducted in R version 4.0.0 [4] using the lqmm [5], nlqmm [6], and aqmm [7] packages.

#### Correlations among Tmax indices

We used the same specifications to correlate the different Tmax indices among them. For terrestrial species, this included the following correlations: Tmaxexp X Tmax prot, Tmaxexp X Tmaxair, Tmaxprot X Tmaxair. For marine fish, we correlated Tmaxfsurf with Tmaxmid depth, both obtained from the Bio-Oracle dataset. Results are shown in the supporting tables set 2 and in Figure 2B.

#### Test of differences in effect sizes across quantile correlations of Tmax-Heat tolerance

To test if tolerance impacts Tmax more strongly when species are more challenged by environmental temperatures, we tested whether the effect size of the quantile correlations would change across quantiles. For mixed models, the effect size can be represented by the slope of linear models^32^, and we used the scale of the non-linear mixed models for fish. For that, we fitted another mixed model (lmer function from lme4 package^33^). Here, quantile was the fixed effect (categorical, three levels), and the grouping variables were: dataset (five levels) and group (thirteen levels). In this fashion, we directed all the available degrees of freedom (from 117 effect size estimates) to test for a general effect of the fixed effect (i.e across taxonomic groups, indices, and datasets), keeping comparisons of effect sizes within the same dataset and taxonomic group. For example, the effect size for the 10^th^ quantile correlation obtained for fish’s Tmax surf-BIORACLE dataset was compared only with the correlations for the 50^th^ and 90^th,^ of the same dataset and animal group. Finally, we calculated statistical significance for the fixed effects only, using the Kenward-Roger method for p-value calculation (LmerTest^34^; and pbkrtest^35^, packages). Full results are given in the supporting tables 2 (2D).

Scripts for all analyses are available under request.

## Acknowledgements

Supporting fellowships for ACG: (PNPD-CNPq, process: 0001; Fapesp-BEPE, 015-003, MSCAH2020: 897901), UIDB/04292/2020; UIDB/04326/2020), FCT/FEDER (28014 02/SAICT/2017); MTR (FAPESP 2011/50146-6 and CNPq)

## Author contributions

Conception: ACG/MTM, data access and fieldwork: ACG/RJ/MTR/MH, Analyses: ACG/MG, Writing: ACG/MTM, Review: ACG/MTR/MG/CV/MC.

## Additional information

**Supplementary Information** is available for this paper.

Correspondence and requests for materials should be addressed to AC.

**Extended data figure/table legends.**

